# Study of the anti-inflammatory and anti-apoptosis effect of the traditional Mongolian Hohgardi-9 in acute lung injury

**DOI:** 10.1101/2022.07.14.500152

**Authors:** Aodeng Qimuge, Bilige Bilige, Wuhan Qimuge, Siqin Siqin, Hugelile Hang, Temuqile temuqile, Shana Chen, Huricha Baigued, Changshan Wang, Tegexi Baiyin, Dezhi Yang

**Affiliations:** Pharmaceutical Laboratory, Inner Mongolia International Mongolian Hospital, Hohhot 010065, China; Inner Mongolia Traditional Chinese&Mongolian Medical Research Institute, Hohhot 010017, China; Institute of Mongolian Medicinal Chemistry, School of Chemistry and Chemical Engineering, Inner Mongolia University, Hohhot 010021, China; School of Life Sciences, Inner Mongolia University, Hohhot 010021, China

**Author notes:** Correspondence to: Tegexi Baiyin: Pharmaceutical Laboratory, Inner Mongolia International Mongolian Hospital, Hohhot, Inner Mongolia 010065, P. R. China., Dezhi Yang: Pharmaceutical Laboratory, Inner Mongolia International Mongolian Hospital, Hohhot, Inner Mongolia 010065, P. R. China. Aodeng Qimuge and Bilige Bilige have contributed equally to this work.

## Abstract

**Objective:** To explore the main active components of Hohgardi-9 and its mechanism treating in ALI.

**Methods:** Through searching the TCMSP database, we obtained the main components and action targets of Hohgardi-9, and the targets related to ALI were analyzed as the possible targets of Hohgardi-9. Then, the compound target network was constructed using Cytoscape software and obtained the key compounds of Hohgardi-9 acting on ALI. The blood entering components of Hohgardi-9 were analyzed by metabonomics. Using a string database to investigate the interaction between proteins of possible targets of Hohgardi-9, Gene Ontology (GO) function annotation and Tokyo Encyclopedia of the genome (KEGG) pathway enrichment analysis were carried out at the same time to predict its mechanism. Finally, the ALI rat model verified the pharmacodynamic effects and key targets of Huhgridi-9.

**Results:** The network pharmacology and blood component analysis results showed that 27 potentially active components such as quercetin, herbacetin, izoteolin, and columbinetin acetate were the major functional components in Hohgardi-9. Those might act on NF kappa B signalling pathway, toll-like receptor signalling pathway, and TNF signalling pathway through key targets such as RELA (p65), TLR4, etc. In vivo experiments showed that Hohgardi-9 significantly improved lung tissue injury and pulmonary edema in ALI rats. At the same time, the Hohgardi-9 intervention could significantly reduce the mRNA expression levels of TRL4, TNFa, IL-1 β, and ICAM1 in ALI rats.

**Conclusion:** Hohgardi-9 revealed ALI through the inhibiting inflammatory factor apoptosis-related gene expression.

## 1 Introduction

Acute lung injury (ALI) is a kind of respiratory disease with high morbidity and mortality, and its clinical manifestations include refractory hypoxemia, acute respiratory distress and non-cardiogenic pulmonary edema. Its main pathogenesis is the release and activation of inflammatory factors in the lungs induced by various etiologies, resulting in damage to alveolar epithelial cells, increased alveolar permeability, and eventually pulmonary edema and even respiratory failure[1, 2]. Therefore, controlling the inflammatory response can alleviate the progression of ALI to a certain extent. ALI is a common pathological reaction of many diseases, such as SARS, Influenza A and COVID-19. Their pathogenesis and cause of death are closely related to the occurrence and development of ALI[3-5]. From the therapeutic effect of traditional medicine for COVID-19, traditional medicine reveals its unique advantages and functions in improving the recovery rate, inhibiting the transition to severe disease and reducing sequelae, etc., i.e., from mild symptoms, moderate symptoms and severe symptoms to rehabilitation periods[6, 7].

According to traditional Mongolian medicine, COVID-19, a “plague fever” is one acute infectious disease caused by virus infection[7]. In the Traditional Mongolian medicine theory, a disease causes weak body and poor immunity by disturbing the balance of the three elements(Heyi, Xila and Badagan) of life, invading the body by “Nian” (pathogenic microorganisms); it was fighting with Heyi, Xila and Badagan, and finally damaging the organs[7, 8].

Hohgardi-9, also known as Qinggan-9 flavour pills, is a classic prescription for the traditional treatment of influenza and other plagues in Traditional Mongolian medicine. It is composed of 9 herbs, including Zhicaowu (*Aconitum carmichaelii*), Hezi (*Terminalia chebula*), Fanbaicao (*Potentilla discolour*), Tumuxiang (*Inula helenium*), Louhu *(Stemmacantha uniflora*), Heiyunxiang (resin of *Commiphora Mukul*), Quanshen (*Bistorta officinalis*), Beishashen (*Adenophora Borealis*) and Huhuanglian (*Neopicrorhiza scrophulariiflora*)[9].It eliminates viscosity, relieves heat and treats coughs, and is mainly used for treating plague fever, cold coughs and sore throat[10].

Here, the network pharmacology and in vivo experiments were ues to uncover the mechanism of Hohgardi-9 in the treatment of lipopolysaccharide (LPS) -induced rat acute lung injury model.

## 2 Methods

### 2.1 Active components of Hohgaridi 9

Based on the Traditional Chinese Medicine Systems Pharmacology Database (TCMSP) and using a literal search for the chemical composition of Beishashen, Caowu, Fanbaicao, Hezi, Huhuanglian, Louhuhua, Tumuxiang, Quanshen and Heiyunxian, oral bioavailability and drug-likeness were used as screening conditions to choose compounds with higher activity. A total of 78 compounds were obtained, including eight cayenne compounds, 8 from Hezi, 5 from Tumuxiang, 5 from Louluhua, 24 from Heiyunxiang, 6 from Quanshen, 8 from Beishashen, 5 from Fanbaicao and 10 from Huhuanglian.

### 2.2 Active ingredient target proteins and disease-associated target proteins

Targets of candidate compounds with a probability >0 are involved through component prediction using the Swiss Target Prediction database[11] (http://www.swisstargetprediction.ch/). All targets are screened for the species of “human” and removed for duplicates using the protein database[12] (Uniprot Database,http://www.uniprot.org/), and then related target information to candidate compounds can be obtained. Among 78 candidate compounds, 72 candidate compounds matched human targets in the database. Six candidate compounds failed to match the relevant targets enter “acute lung injury” into the DisGeNET database (https://www.disgenet.org/)[13], NCBI Genetic Database (https://www.ncbi.nlm.nih.gov/) and GeneCard database https://www.genecards.org/respectively and search for drug targets associated with the treatment for acute lung injury to obtain known disease-related target data from these three databases and create disease target datasets as a backup[14][15].

### 2.3 Build a network diagram

The Bioinformatics & Evolutionary Genomics, an online software drawing tool platform. (http://bioinformatics.psb.Ugent.be/webtools/Venn//) was used to map the potential chemical targets of Hohgardi-9 Flavors Pills with disease-related targets to construct a Venn diagram. The common targets with an intersection may be the key targets for treating acute lung injury using Hohgardi-9. To better understand the complex relationships between components, diseases, and corresponding targets, Cytoscape 3.8.0 software was used to construct a composition-target network diagram, visualizing the relationship between compounds and target protein.

A protein-protein interaction (PPI) network model was constructed by placing common target proteins on the STRING database (https://string-db.org/cgi/input.pl) to study the links between target proteins and input data into Cytoscape3.6.2 to obtain a PPI visualization than can give a visual analysis of key proteins[16].

### 2.4 GO and KEGG Pathway Enrichment Analysis

The biological process (BP) enrichment of GO was conducted on common targets of drug diseases. The String database was introduced to screen out the items with P whose corrected value is lower than 0.05. Meanwhile, the KEGG pathway enrichment analysis was performed on common targets of drug diseases; the String database was introduced to screen out the items whose corrected value is lower than 0.05.

### 2. 5 A analysis of blood components of Hohgardi-9

#### 2.5.1 Administration

After one week of adaptive feeding, all male SD rats were divided into the Blank Control Group and the Hohgardi-9 Group. In the Hohgardi-9 Group, all rats were administrated five times in total, twice a day in the first two days, fasted on the second night, and drawn for blood one hour after administration on the third day.

#### 2. 5.2 The Collection and processing of plasma samples

Centrifugation was performed at 3 000 g/ min for 10 mins. The supernatant was taken to obtain a plasma sample. The operation was performed on the ice. 50 μL of plasma sample was transferred into 1.5 mL EP tube, then 200 μ L of cold methanol (with internal standard) was added, with vortex oscillation for 2 mins and let stand at low temperature for 10 mins. 14,000 g was centrifuged for 15 min at 4 °C. 200 μL of supernatant was drained into a new EP tube. After it was centrifuged and concentrated at low temperature, the sample was resold with 100 μL of 20% methanol/aqueous solution for positive and negative ion pattern analysis.

#### 2.5.3 LC-MS Analysis

##### Mass Spectrometry Conditions

LEVEL 1 full scan using mass spectrometry+DDA Secondary subion ion scanning -Positive ion mode: HESI-Positive, heated electrospray ion source model, is used. Leve 1 Full Scan and DDA Secondary subion ion scanning were adopted. Spray Voltage (kV): +3.8; Capillary temperature (°C): 320; Aux gas heater temperature (°C): 350; Sheath gas flow rate (Arb): 35; Aux gas flow rate.

(Arb): 8; S-lens RF level: 50; Mass range (m/z): 100-1200, Full ms resolution: 70000; MS/MS resolution:17500; TopN: 5; NCE/stepped NCE: 20,40 _∘_ LEVEL 1 full scan using mass spectrometry+ DDA Secondary subion ion scanning -Negative ion mode: HESI-Negative mode, heated electrospray ion source, is adopted, Level-1 full scan was used and +DDA Secondary subion ion scanning Spray Voltage (kV): -3.0; Capillary temperature (°C): 320; Aux gas heater temperature (°C): 350; Sheath gas flow rate (Arb): 35; Aux gas flow rate (Arb): 8; S-lens RF level: 50; Mass range (m/z): 100-1200, Full ms resolution: 70000; MS/MS resolution:17500; TopN: 5; NCE/stepped NCE: 20.

Chromatographic conditions: Positive ion mode: Chromatographic column: Waters Corporation USA, BEH C8 Column: (1.7 μm, 2.1 ×100 mm), Column Temperature: 50°C, Injection volume:10 μL, A Spectrum: Water (0.1% formic acid), B Spectrum: Acetonitrile (0.1% formic acid), Flow Velocity: 0.35 ml/min, Gradient elution procedure: 0-1 min, 5% (B); 1-24 min, 5%-100% (B); 24.1-27.5 min, 100% (B); 27.6-30 min, 5% (B). Negative ion mode: Chromatographic column: Waters Corporation USA, HSS T3 Column: (1.8 μm,2.1×100 mm), Column Temperature: 50°C, Injection volume:10 μL, A Spectrum: Water (6.5 mM ammonium bicarbonate), B Spectrum: 95% Methanol water (6.5 mM ammonium bicarbonate), Flow Velocity: 0.35 ml/min, Gradient elution procedure: 0-1min, 5% (B); 1-18 min, 5%-100% (B); 18.1-22 min, 100% (B); 22.1-25 min, 5% (B).

### 2.6 In vivo experimental validation

#### 2.6.1 To establish a model of a rat with acute lung injury

30 SD rats were taken, randomly grouped, 6 per group. After one week of adaptive feeding, a blank group, LSP model group, TCM Positive Control (Lianhua Qingwen, 4g/kg), Positive Control (aspirin, 0.1g/kg), the Hohgardi-9 Group (2 g/kg, five times the dosage of humans), High Dosage (medium dosage, ten times the dosage of human), the Hohgardi-9 Group, Low Dosage (Low dosage, five times the dosage of human), All rats were administrated one time per day for seven consecutive days. The same volume of normal saline was given to the Blak Group and the Model Group. Except for the blank group, LPS solution (5 mg/kg) was intraperitoneally injected one hour after administration on the seventh day to build an acute lung injury model in rats. Six hours after modelling, samples were taken and executed.

#### 2.6.2 Dying/histological observation of the lungs

Lung tissue was fixed in 10% formaldehyde solution, dehydrated with ethyl alcohol, embedded with paraffin and sliced, with the thickness of 5 mm, dyed with HE and pathological changes in lung tissues were observed.

#### 2.6.3 Wet-to-dry (W/D) lung weight ratio

To assess pulmonary edema, the left lung was collected, briefly rinsed in PBS, aspirated, and weighed to obtain wet weight. The lung tissue is then placed in a thermostatic oven at 68 °C for 48h to obtain dry weight, and the ratio of the wet lung to dry lung was calculated to determine the degree of pulmonary edema.

#### 2.6.4 Real-time Quantitative Fluorescence PCR Experiment

Total RNA was extracted from lung tissues using Trizol, and cDNA was synthesized using Revert Aid First Strand cDNA Synthesis Kit. The templates were amplified by real-time PCR. The reaction procedure is 94°C 30s, 94°C 5s and 60°C 34s (45 cycles). With β-actin as an internal reference, the Ct values obtained were treated by the 2^-CT^ method. Primers used for reverse transcription are shown in Table 1.

**Table 1:**
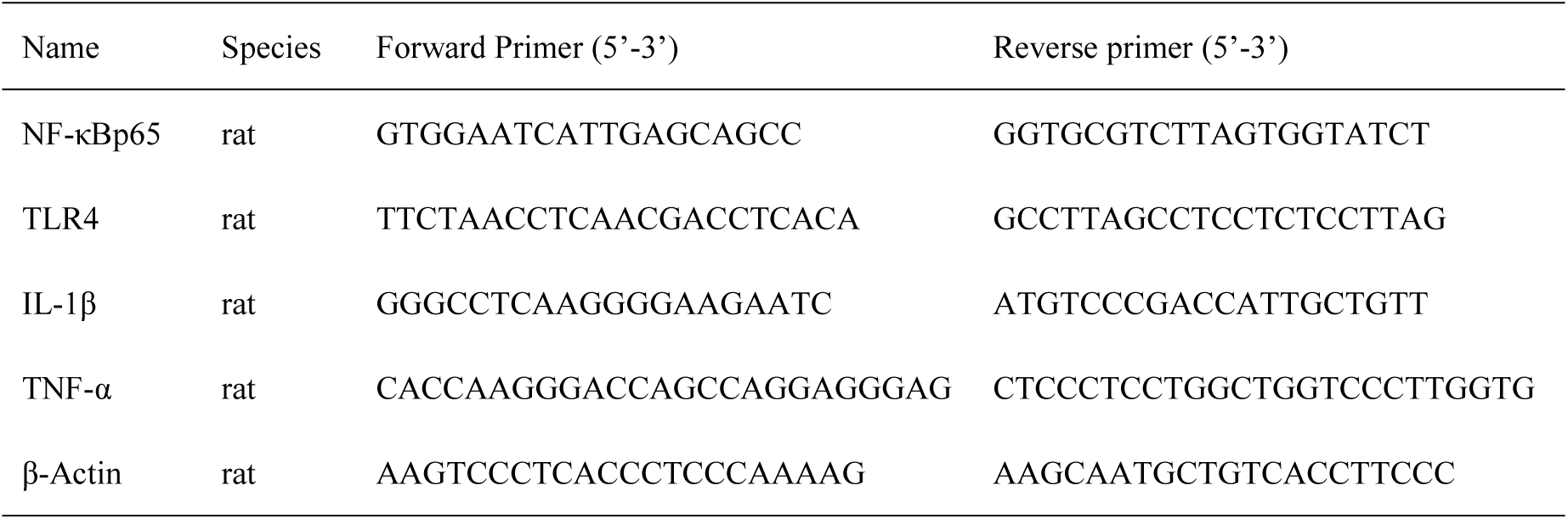
Primers of target genes.

### 2.7 Statistical analysis

Data are presented as “mean ± standard deviation” and SPSS 20.0 was used for statistical analysis.

## 3 Results

### 3.1 Component-target network diagram analysis

Venn diagrams were drawn using 2005 disease-related targets and 848 potential targets of Hohgardi-9. After intersection, 379 common targets were obtained. These are the significant targets of Hohgardi-9 in acute lung injury as shown in Table 1. The relationship between candidate compounds and common targets was constructed into a compound-target protein network (see Figure 2). This network has 442 nodes(including 63 compound nodes and 379 target protein nodes) and 3738 edges, and the average number of binding compounds per target protein was 4.93; 145 targets (38.2%) correspond to more than five small molecules in the target proteins. Each candidate compound acts on an average of 29.7 target proteins. It shows that Hohgardi-9 plays an integral role through multiple components and multiple targets. In candidate compounds, quercetin in the rhododendron has the highest degree value (degree value 64); the following was n-Feruloyltyramine (degree value 63) in Heiyunxiang and kaempferol in Fanbaicao (degree value 62). The degree values of herbacetin in Huhuanglian, izoteolin in Caowu, cheilanthifoline in Hezi, cnidilin in Beishashsen and ellagic acid in Quanshen were all higher than the average value of 29.9. These compounds with higher degree value may be the key compounds for Hohgardi-9 to play a role in treating ALI, as shown in Table 2.

**Fig.1.**
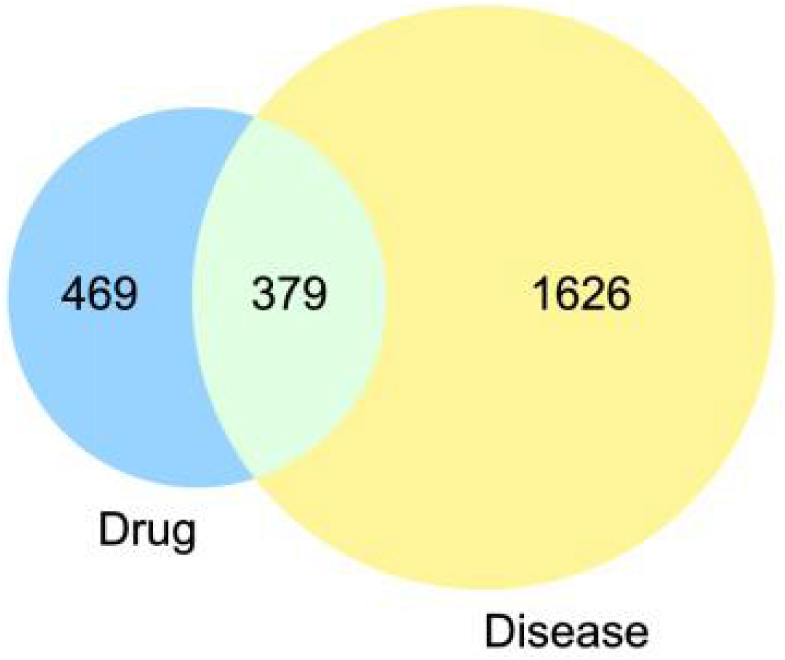
Venn ’s diagram of Acute lung injury targets and Hohgardi-9 component targets

**Fig.2.**
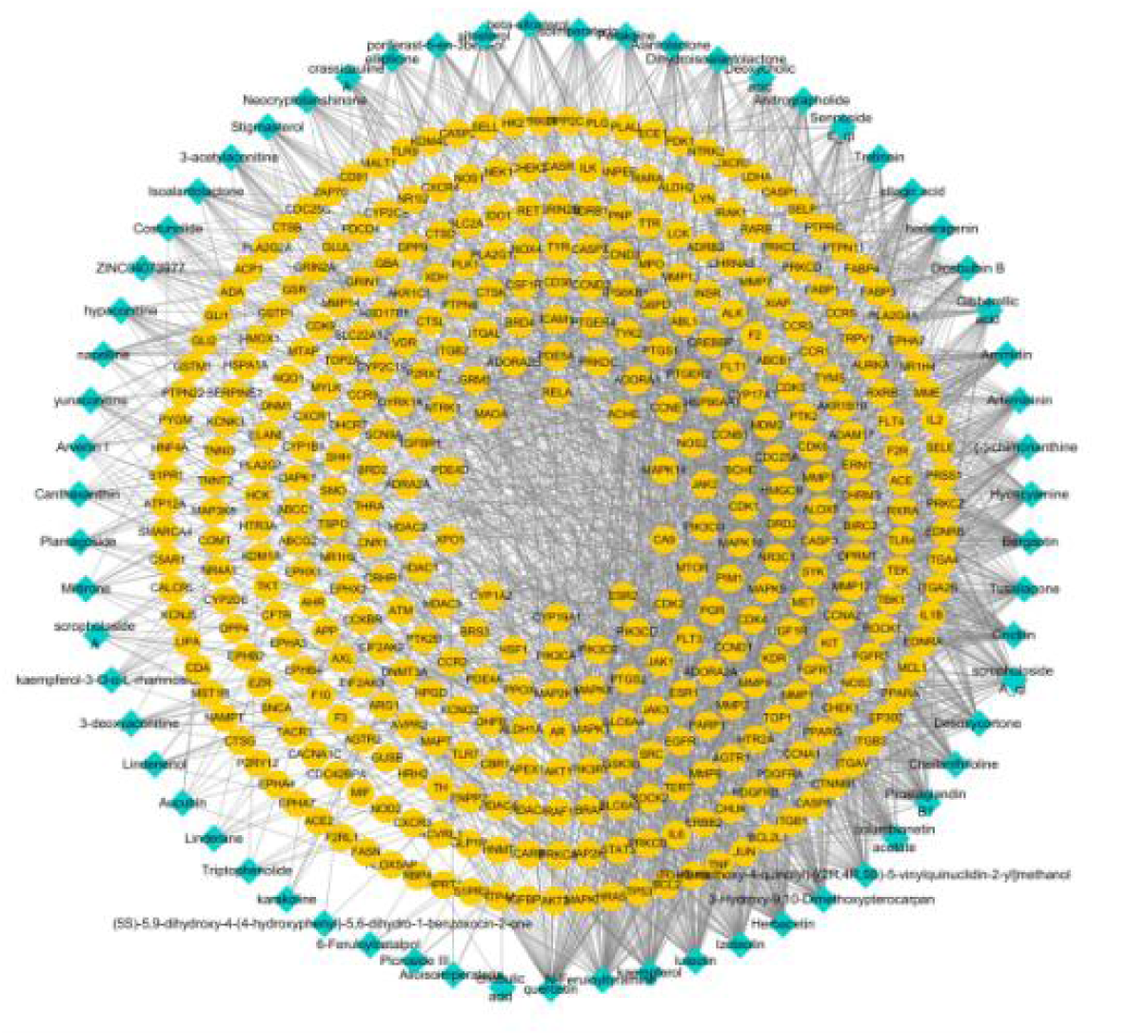
Compound-potential target network (green diamonds represent compounds contained in Hohgardi-9; yellow circle represent compound targets)

**Table 2.**
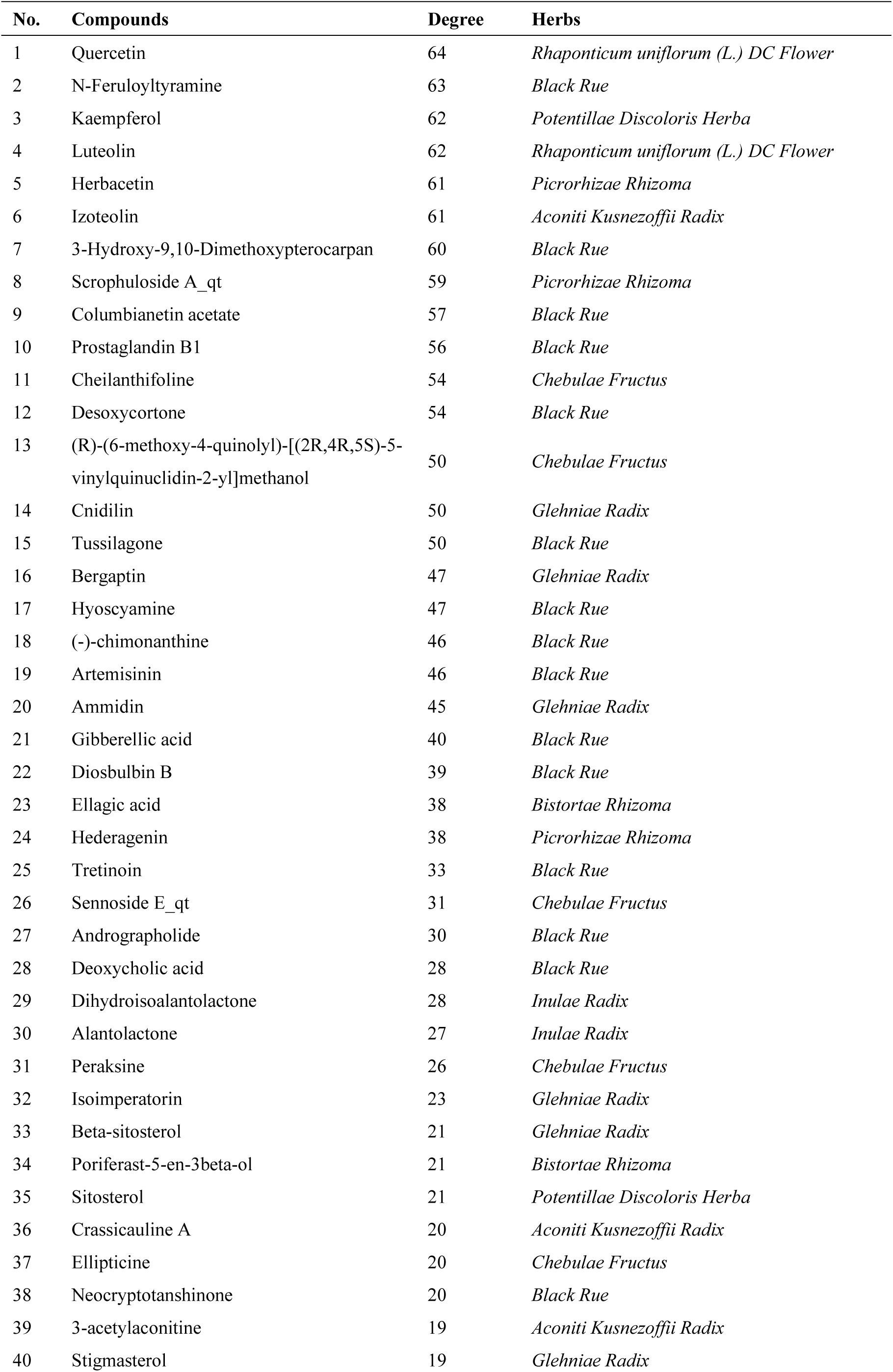
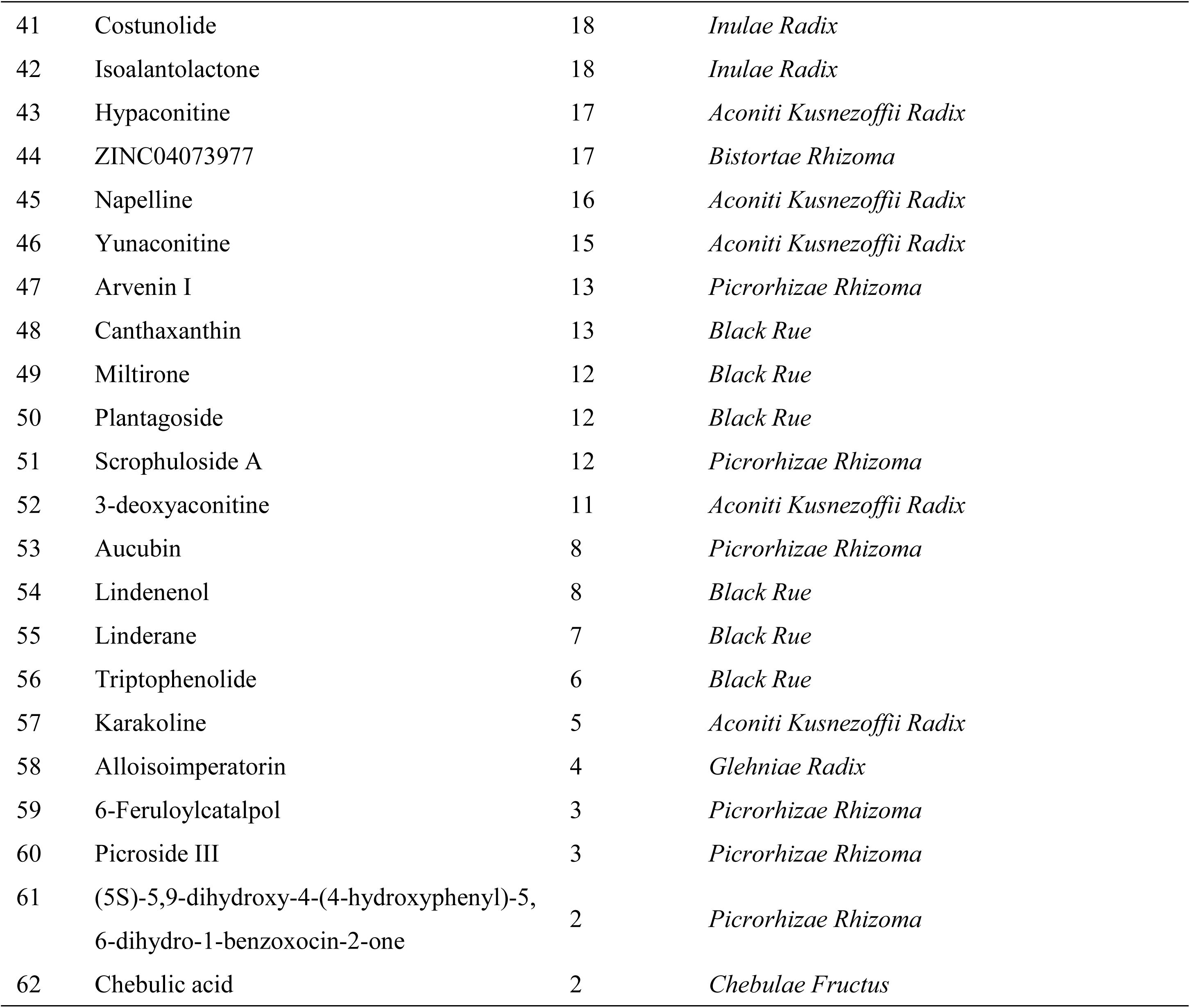
Information of ingredients of Hohgardi-9

### 3.2 Protein Interaction (PPI) Network Diagram Analysis

To better explore how Hohgardi-9 works, 379 common targets were input into the STRING data platform, and a PPI action network was obtained by analysis. A total of 379 interaction nodes and 2013 interaction relationships were received, with an average degree value of 10.6, as shown in Figure 3. In graphical results, the size of nodes represents its degree value. The larger and darker node with a higher corresponding degree value was the more important in the predicted disease. STAT3, MAPK1, AKT1, JUN, RELA, TNF, TLR4, and IL-1 β show a remarkable effect in this result, and these are key targets for inflammatory pathways and may be one of the targets in which Hohgardi-9 plays a therapeutic role.

**Fig.3.**
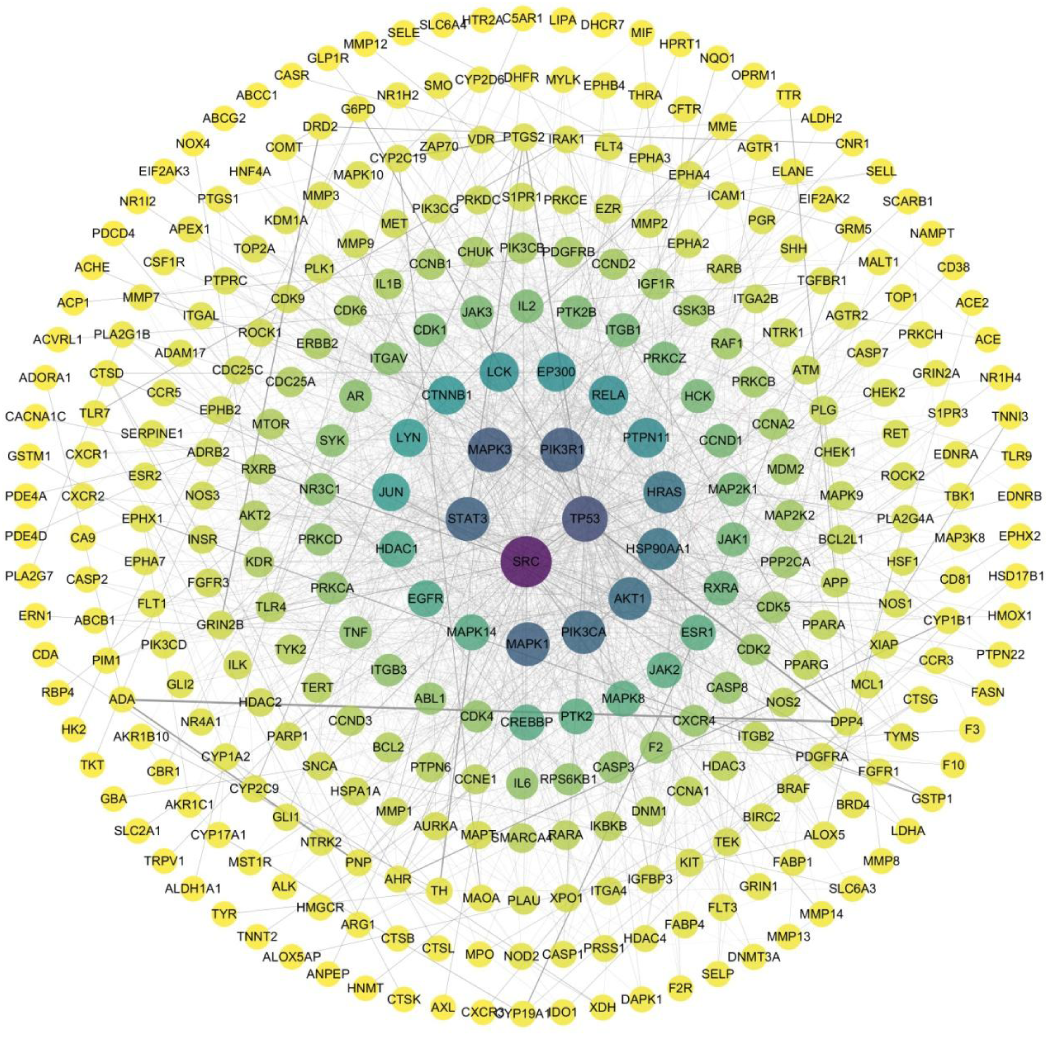
Hohgardi-9 targets’ PPI network (the target gene are sorted according to degree)

### 3.3 GO-BP Enrichment Analysis and KEGG Pathway Enrichment Analysis

For further exploration of the potential mechanism of action of Hohgardi-9, GO-BP enrichment analysis and KEGG pathway enrichment analysis was conducted. In key target proteins, items whose corrected values are lower than 0.05 were screened out using GO-BP enrichment analysis. 101 biological processes were collected in GO-BP, including inflammatory response, response to lipopolysaccharide, response to hypoxia, viral entry into the host cell, apoptotic process, positive regulation of NF-kappa B transcription factor activity, positive regulation of tumour necrosis factor production, response to interleukin-1, as shown in figure 4. There are 122 pathways obtained from KEGG pathway enrichment analysis(P < 0.05), including PI3K-Akt signalling pathway, Apoptosis, HIF-1 signalling pathway, TNF signalling pathway, Toll-like receptor signalling pathway, Coronavirus disease-COVID-19 and NF-kappa B signalling pathway, as shown in figure 5.

**Fig.4.**
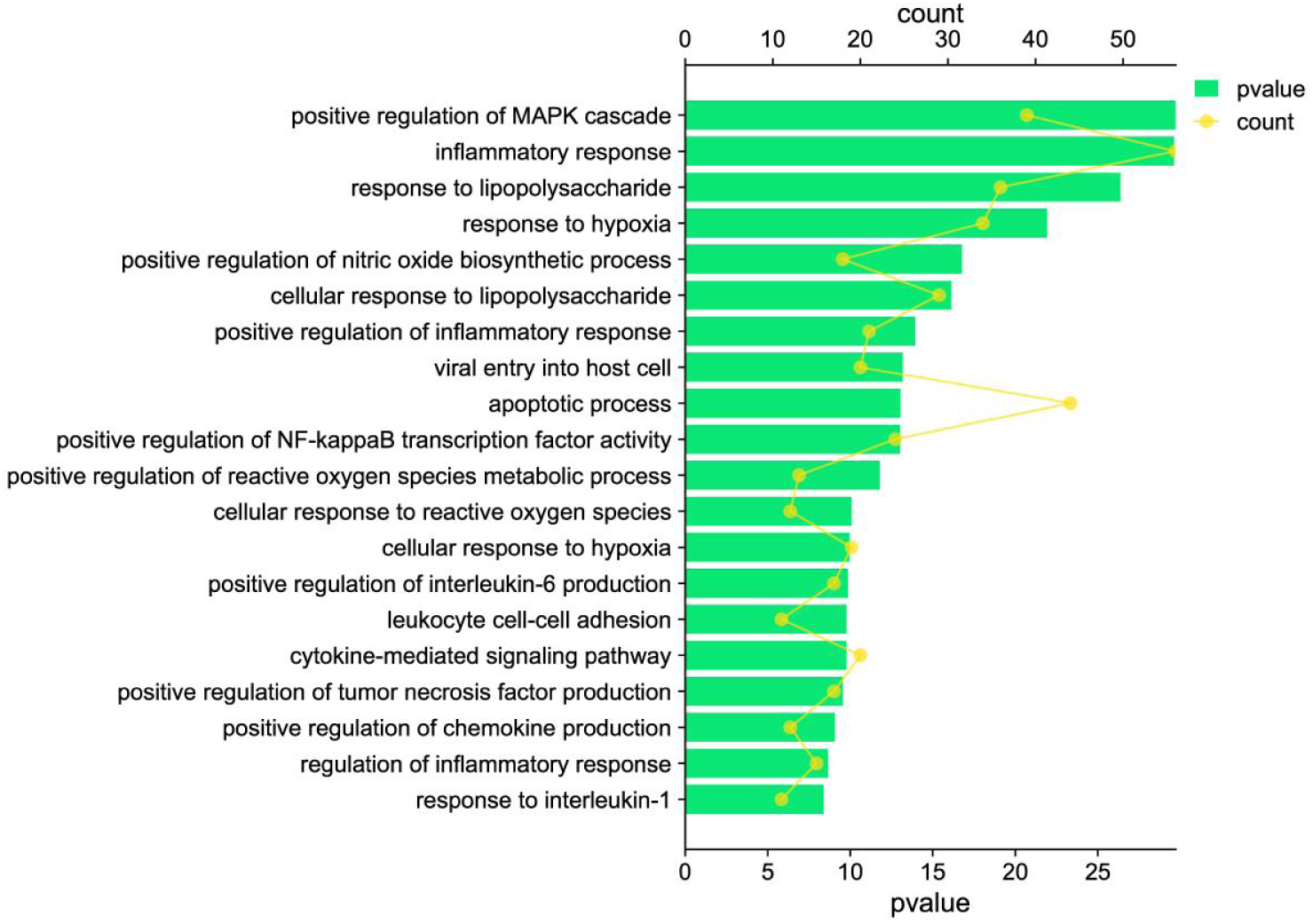
GO enrichment analysis for targets Hohgardi-9

**Fig.5.**
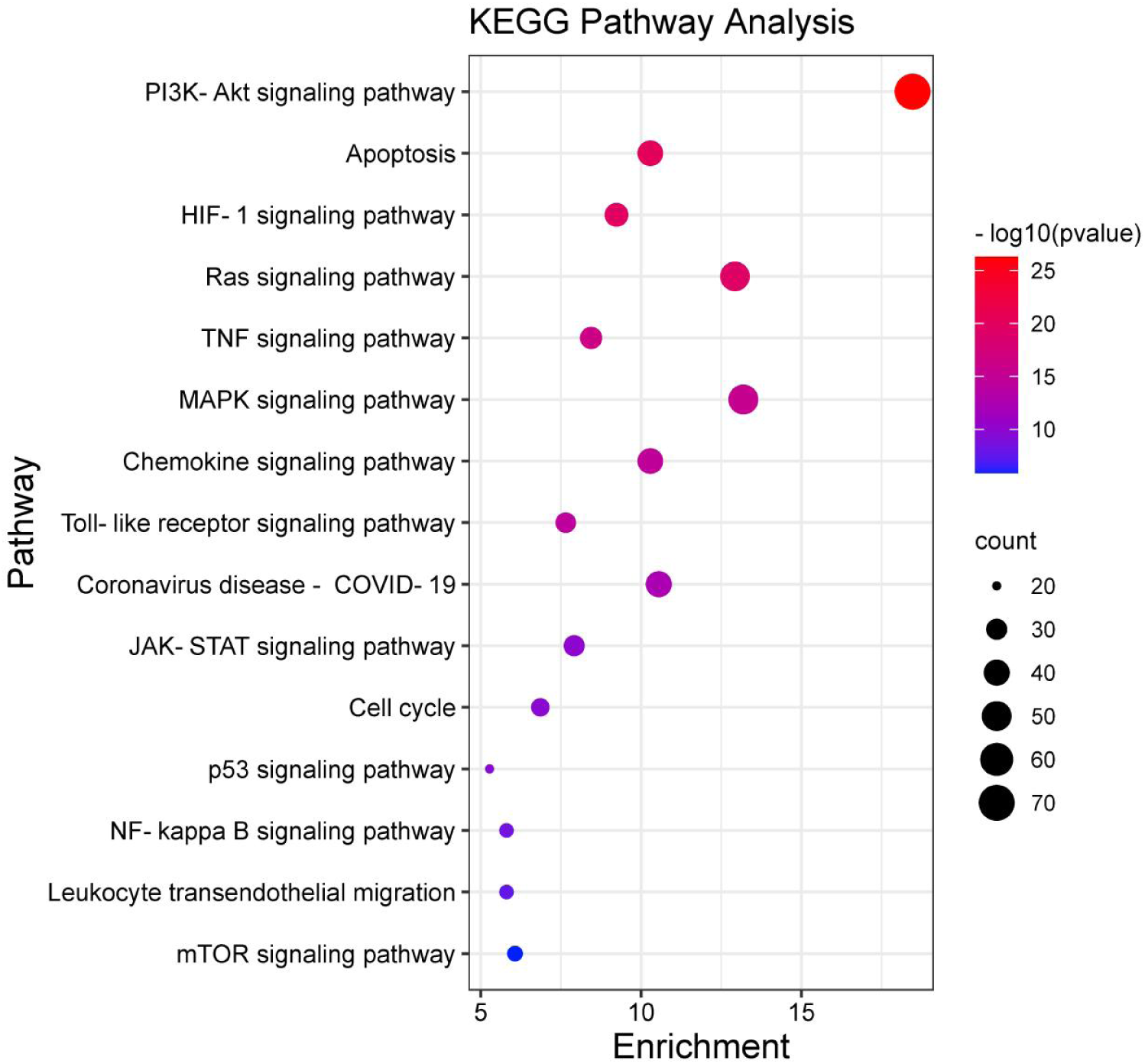
KEGG analysis for the major targets of Hohgardi-9

### 3.4 Blood components of Hohgardi-9

After oral admission of Hohgardi-9, using high-resolution mass spectrometry for the blood drug test, 27 compounds were identified in rat plasma derived from network pharmacologic prediction, as shown in Table 3.

**Table 3.** Identification of the chemical composition of Hohgardi-9 detected in rat plasma.

| No. | R.T. (min) | Mean measured mass (Da) | Elemental composition | Ion mode | Compounds | Herbs |
| --- | --- | --- | --- | --- | --- | --- |
| 1 | 6.11 | 302.0426 | C<sub>15</sub>H<sub>10</sub>O<sub>7</sub> | [M+H]<sup>+</sup> | Quercetin | *Rhaponticum uniflorum (L.) DC Flower* |
| 2 | 13.04 | 302.0426 | C<sub>15</sub>H<sub>10</sub>O<sub>7</sub> | [M+H]<sup>+</sup> | Herbacetin | *Picrorhizae Rhizoma* |
| 3 | 5.30 | 327.1470 | C<sub>19</sub>H<sub>21</sub>NO<sub>4</sub> | [M+H]<sup>+</sup> | Izoteolin | *Aconiti Kusnezoffii Radix* |
| 4 | 11.29 | 288.0997 | C<sub>16</sub>H<sub>16</sub>O<sub>5</sub> | [M+H]<sup>+</sup> | Columbianetin acetate | *Black Rue* |
| 5 | 15.85 | 330.2195 | C<sub>21</sub>H<sub>30</sub>O<sub>3</sub> | [M+H]<sup>+</sup> | Desoxycortone | *Black Rue* |
| 6 | 6.3 | 300.0998 | C<sub>17</sub>H<sub>16</sub>O<sub>5</sub> | [M+H]<sup>+</sup> | Cnidilin | *Glehniae Radix* |
| 7 | 14.77/14.91 | 390.2406 | C<sub>23</sub>H<sub>34</sub>O<sub>5</sub> | [M+H]<sup>+</sup> | Tussilagone | *Black Rue* |
| 8 | 5.22 | 344.1260 | C<sub>19</sub>H<sub>20</sub>O<sub>6</sub> | [M+H]<sup>+</sup> | Diosbulbin B | *Black Rue* |
| 9 | 5.50 | 302.0063 | C<sub>14</sub>H<sub>6</sub>O<sub>8</sub> | [M+H]<sup>+</sup> | Ellagic acid | *Bistortae Rhizoma* |
| 10 | 15.12 | 300.2089 | C<sub>20</sub>H<sub>28</sub>O<sub>2</sub> | [M+H]<sup>+</sup> | Tretinoin | *Black Rue* |
| 11 | 11.97 | 350.2093 | C<sub>20</sub>H<sub>30</sub>O<sub>5</sub> | [M+H]<sup>+</sup> | Andrographolide | *Black Rue* |
| 12 | 14.94 | 310.1681 | C<sub>19</sub>H<sub>22</sub>N<sub>2</sub>O<sub>2</sub> | [M+H]<sup>+</sup> | Peraksine | *Chebulae Fructus* |
| 13 | 13.76 | 643.3356 | C<sub>35</sub>H<sub>49</sub>NO<sub>10</sub> | [M+H]<sup>+</sup> | Crassicauline A | *Aconiti Kusnezoffii Radix* |
| 14 | 11.00/17.52 | 314.1518 | C<sub>19</sub>H<sub>22</sub>O<sub>4</sub> | [M+H]<sup>+</sup> | Neocryptotanshinone | *Black Rue* |
| 15 | 11.79 | 615.3043 | C<sub>33</sub>H<sub>45</sub>NO<sub>10</sub> | [M+H]<sup>+</sup> | Hypaconitine | *Aconiti Kusnezoffii Radix* |
| 16 | 4.61 | 359.2460 | C<sub>22</sub>H<sub>33</sub>NO<sub>3</sub> | [M+H]<sup>+</sup> | Napelline | *Aconiti Kusnezoffii Radix* |
| 17 | 11.70 | 659.3305 | C<sub>35</sub>H<sub>49</sub>NO<sub>11</sub> | [M+H]<sup>+</sup> | Yunaconitine | *Aconiti Kusnezoffii Radix* |
| 18 | 12.10 | 564.3967 | C<sub>40</sub>H<sub>52</sub>O<sub>2</sub> | [M+H]<sup>+</sup> | Canthaxanthin | *Black Rue* |
| 19 | 9.80 | 282.1620 | C<sub>19</sub>H<sub>22</sub>O<sub>2</sub> | [M+H]<sup>+</sup> | Miltirone | *Black Rue* |
| 20 | 7.50 | 474.1526 | C<sub>24</sub>H<sub>26</sub>O<sub>10</sub> | [M+H]<sup>+</sup> | Scrophuloside A | *Picrorhizae Rhizoma* |
| 21 | 12.74 | 629.3199 | C<sub>34</sub>H<sub>47</sub>NO<sub>10</sub> | [M+H]<sup>+</sup> | 3-deoxyaconitine | *Aconiti Kusnezoffii Radix* |
| 22 | 4.01 | 346.1264 | C<sub>15</sub>H<sub>22</sub>O<sub>9</sub> | [M+H]<sup>+</sup> | Aucubin | *Picrorhizae Rhizoma* |
| 23 | 11.06 | 230.1307 | C<sub>15</sub>H<sub>18</sub>O<sub>2</sub> | [M+H]<sup>+</sup> | Lindenenol | *Black Rue* |
| 24 | 14.11 | 312.1725 | C<sub>20</sub>H<sub>24</sub>O<sub>3</sub> | [M+H]<sup>+</sup> | Triptophenolide | *Black Rue* |
| 25 | 4.19 | 377.2566 | C<sub>22</sub>H<sub>35</sub>NO<sub>4</sub> | [M+H]<sup>+</sup> | Karakoline | *Aconiti Kusnezoffii Radix* |
| 26 | 5.92 | 538.1686 | C<sub>25</sub>H<sub>30</sub>O<sub>13</sub> | [M+H]<sup>+</sup> | 6-Feruloylcatalpol | *Picrorhizae Rhizoma* |
| 27 | 1.05 | 356.0379 | C<sub>14</sub>H<sub>12</sub>O<sub>11</sub> | [M+H]<sup>+</sup> | Chebolic acid | *Chebulae Fructus* |

### 3.5 Hohgardi-9’ Effects on lung tissues

The experimental results showed that the rats in the blank group showed no obvious morphological damage. Compared with the blank group, rats in the model group showed significant lung injury reaction, with alveolar wall thickening, septal edema, numerous neutrophils and a few eosinophils infiltration mast-cells around blood vessels, and foam cells and haemorrhage in a few alveolar lumens. The tissue damage was significantly improved due to the effect of Hohgardi-9 and Lianhua Qingwen. Results are shown in Table 6.

### 3.6 Hohgardi-9’s effects on the lung wet-dry ratio induced by lipopolysaccharide

To further verify the modelling success, the pulmonary wet-dry weight ratio (W/D ratio) was quantified to indicate pulmonary edema. As shown in figure 7, compared with the blank group, with LPS stimulation, the lung W/D ratio increases significantly (p < 0.05). However, with medication, these symptoms are significantly relieved(p < 0.05).

**Fig.6.**
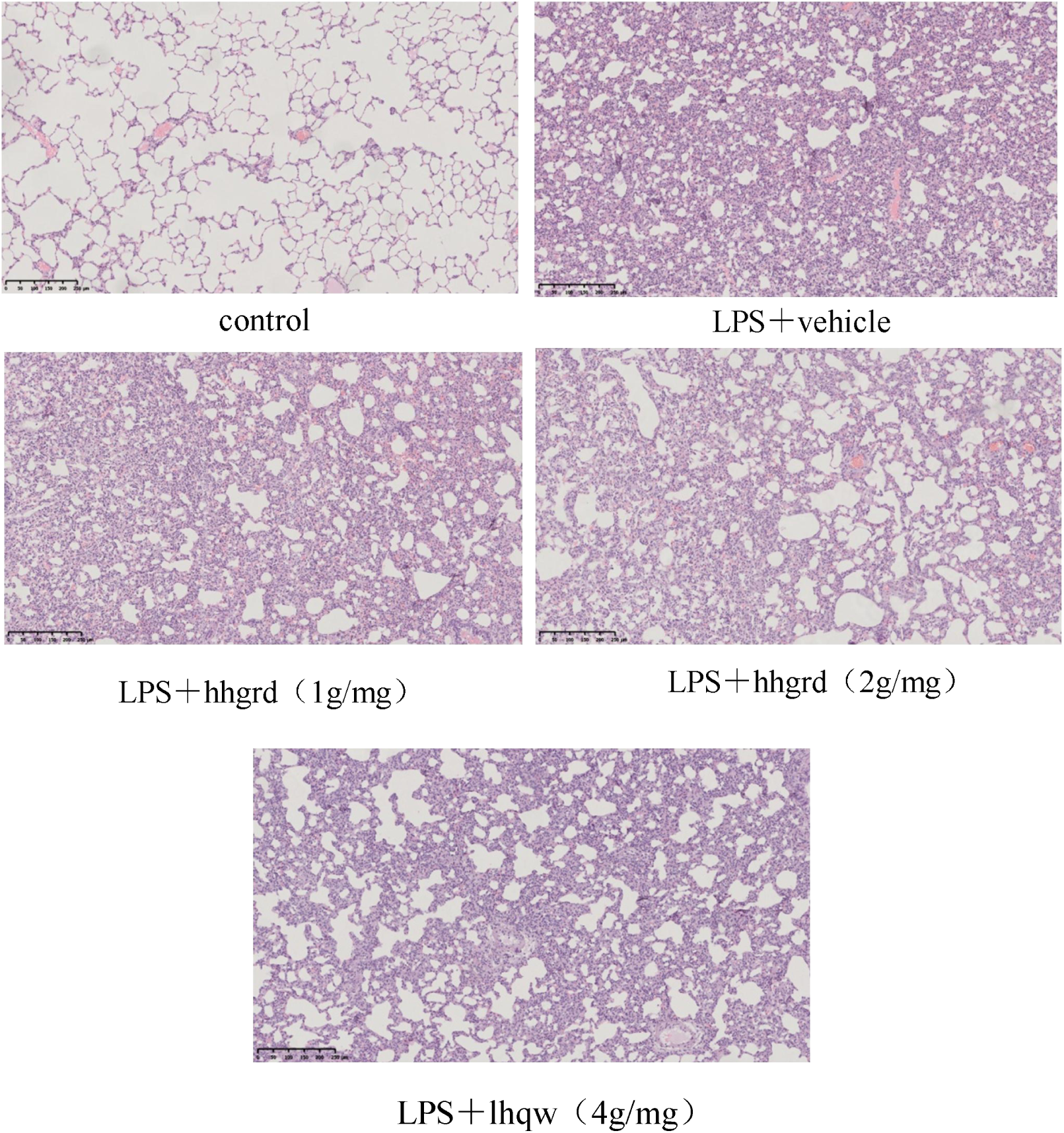
Effects of Hohgardi-9 on pathological changes of lung tissue in acute lung injury mice (HE staining)

**Fig.7.**
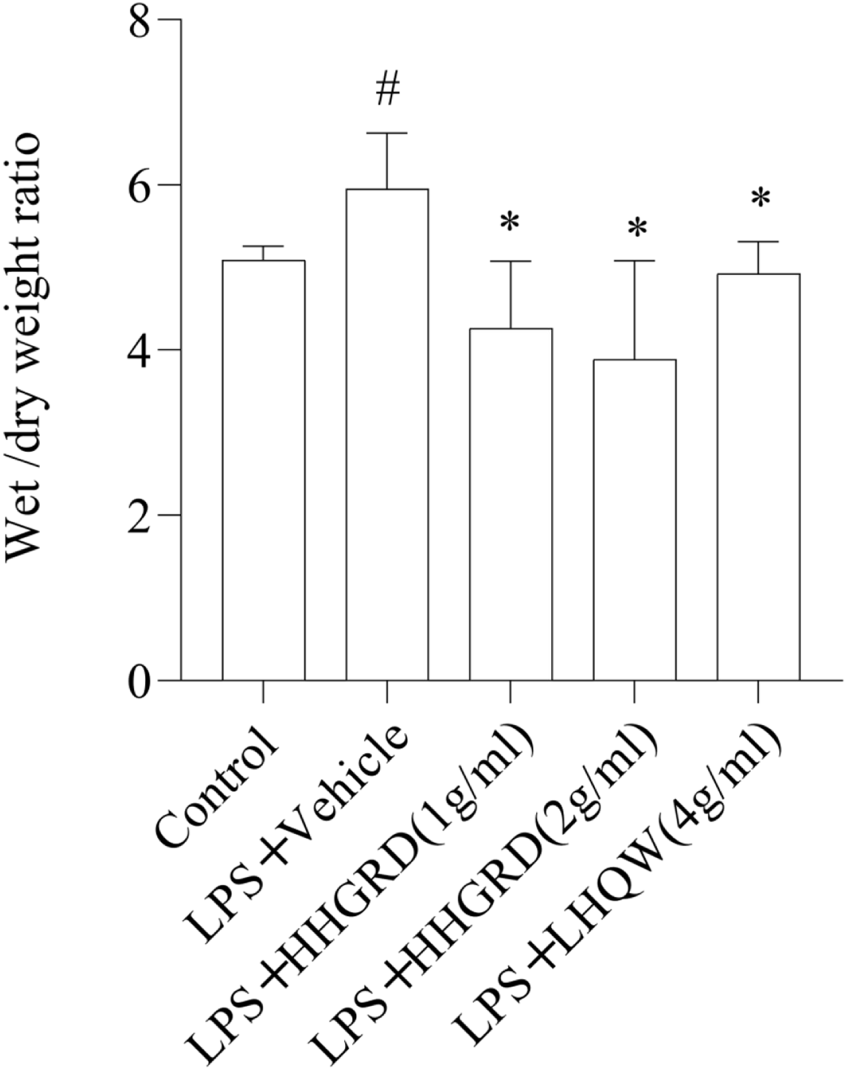
Effects of Hohgardi-9 on LPS induced lung wet-to dry ratio. All the values are stated as mean ± standard deviation (n = 4). ##p < 0.01 and #p < 0.05 versus control group. **p < 0.01 and *p < 0.05 versus LPS group.

### 3.7 Validation of key target mRNA expression with PCR

Based on the network prediction results, 4 key targets, namely TRL4, TNFa, IL-1β and ICAM1, were verified by a real-time quantitative PCR experiment. The results show that compared with the blank group, the mRNA expression of TRL4, TNFa, IL-1β, and ICAM1 in lung tissues in the model group was significantly increased (**P < 0.01). Compared with the model group, The expression of TRL4, TNFa, IL-1β and ICAM1 were significantly decreased by the intervention of Hohgardi-9 (*P < 0.05, **P < 0.01). The results are shown in figure 8. The results showed that the PCR results were consistent with the predicted results.

**Fig.8.**
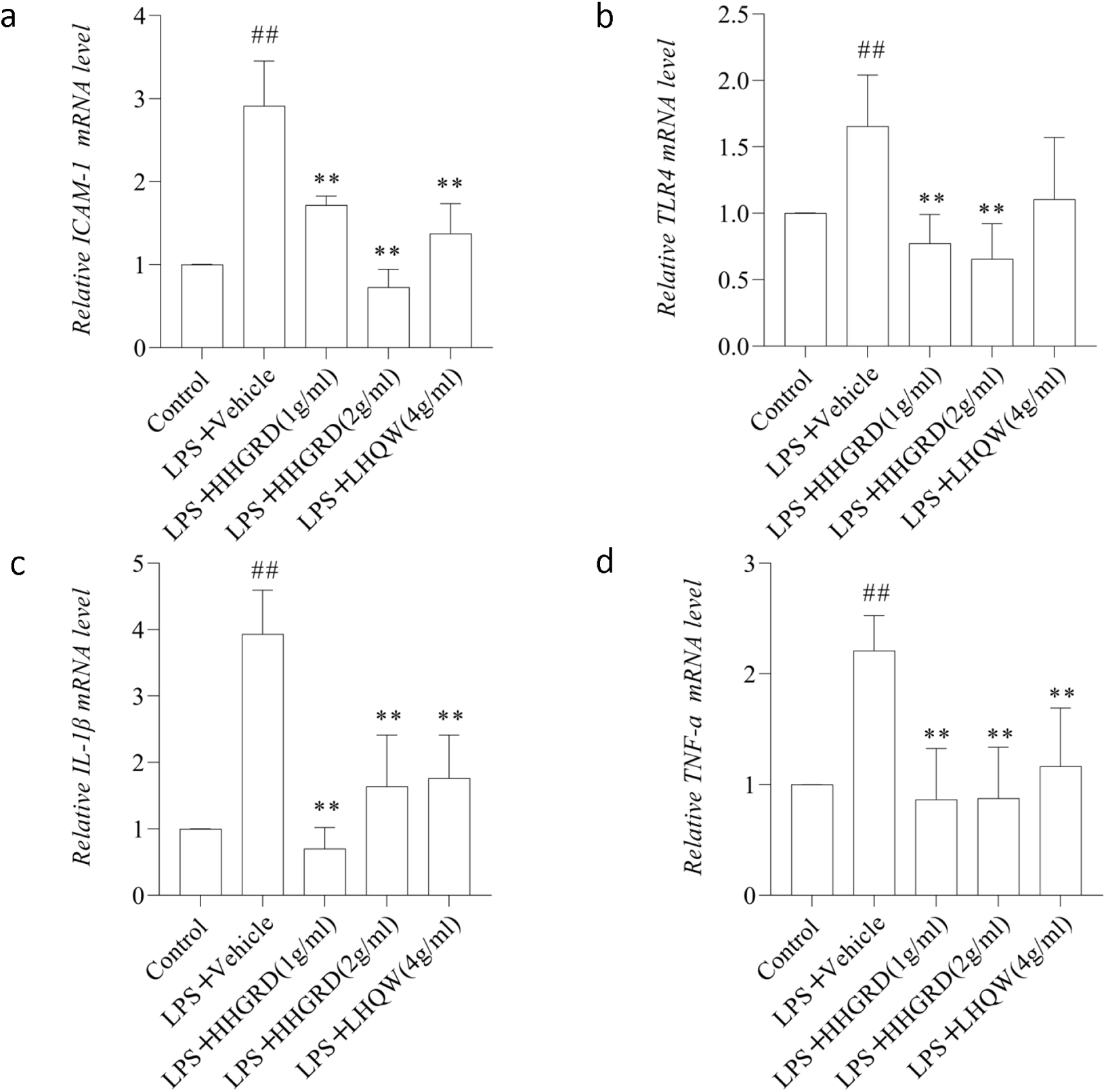
The levels of ICAM-1(a), TLR4(b), IL-1β (c), TNF-a(d) mRNA in lung assessed by Real time-PCR. All the values are stated as mean ± standard deviation (n = 5). ##p < 0.01 and ###p < 0.05 versus control group. **p <0 .01 and ***p <0 .05 versus LPS group.

## 4. Discussion

Acute lung injury (ALI), a common pathological process of many diseases, including COVID-19, is a serious disease with high incidence and fatality rates[17]. In this paper, the mechanism of action of Hohgardi-9 in treating acute lung injury was predicted and analyzed using network pharmacology and the relevant targets, and it was found that an inflammatory response to oxidative stress and apoptosis pathways were involved in this procedure.

The analysis of composition-target network diagrams shows that in candidate compounds, quercetin, columbianetin acetate, kaempferol, herbacetin, izoteolin, cheilanthifoline, cnidilin, and ellagic acid may be the main active components of Hohgardi-9 in treating of ALI. An analysis of blood components of Hohgardi-9 shows among the candidate compounds, 29 compounds were identified as blood constituents, including compounds with higher degree value: quercetin, columbianetin acetate, herbacetin, and columbianetin acetate. izoteolin cnidilin, ellagic acid compounds, etc. For example, quercetin is a polyhydroxyflavonoid compound and has the pharmacological effect of anti-inflammation, anti-bacteria and anti-virus and immunoregulation[18, 19]. Studies demonstrate that quercetin shows a protective role in acute lung injury induced by lipopolysaccharides in rats[20]. This may be related to Hohgardi-9’s effect; quercetin can affect the expression of the antioxidant protein in the NRF-2ARE signalling pathway, down-regulate the expression of NF-κ Bp65 and ICAM-1, and reduce the release of pro-inflammatory factors[21, 22]. Ellagic acid is a natural polyphenol active ingredient with antioxidant and anti-inflammatory properties. It is reported that ellagic acid can inhibit dermatophagoides pteronyssinus-induced inflammatory response of human bronchial epithelial cells by regulating STAT3 and NF-κB signalling pathways[23].Studies have also indicated that ellagic acid can inhibit the inflammatory response and oxidative stress by inhibiting activation of the NF-κ B/COX-2 inflammatory pathway and have a significant anti-inflammatory effect[24]. Columbianetin, a coumarin chemical compound, has a variety of biological activities, including anti-inflammation, antioxidant, anti-tumour and anti-bacteria[25, 26]. Studies show that it has good anti-inflammatory activity in vitro. It can inhibit the release of a variety of cytokines and nitric oxide by regulating MAPKs and NF-κ B signals and thus improve inflammatory response[27]. Besides serving a good protective role in the ALI mouse model, columbianetin also showed an anti-inflammatory and protective role by inhibiting the activation of NOD1/NF-κ B pathways[27].

According to the PPI network, there are multiple associations between targets. Also, the higher the degree, the better the target’s potential therapeutic effect. Here, TP53, MAPK3, STAT3, MAPK1, RELA, JUN, MAPK14, TNF, CASP3, TLR4, IL-1β, and BCL2 are important targets for inflammatory and apoptosis pathways and the key therapeutic targets for Hohgardi-9. TP53, CASP3, and BCL2 play an important role in the regulation of apoptosis signal transduction, and the ALI mechanism has been proved to be closely related to apoptosis[28]. Inhibiting the activation of TLR-4/NF-κB signal pathways can reduce the secretion and expression of inflammatory cytokines, such as TNF-α and IL-1β and reduce the inflammatory cascade reaction and thus reduce inflammation’s damage to the lung[29, 30]. The GO and KEGG pathways analysis shows that the NF-kappa B signalling pathway, TNF signalling pathway, MAPK signalling pathway, apoptosis, p53 signalling pathway, HIF-1 signalling pathway and Coronavirus disease - COVID-19 were the key targets signal pathways. Based on those we know, Hohgardi-9 can treat ALI by reducing the inflammatory response and oxidative stress, inhibiting Apoptosis and fighting the virus. Animal experiments confirm that Hohgardi-9 reduce rats’ lung injury, relieves pulmonary edema and decreases the expression of inflammation-associated genes mRNA.

In conclusion, this study explores active ingredients, targets and related pathways in treating ALI using Hohgardi-9 through network pharmacology. It verifies its efficacy and the regulation mechanism of Hohgardi-9 through animal experiments and molecular biological methods. This study shows that Hohgardi-9 successfully relieves LPS-induced ALI by inhibiting inflammatory factors and apoptosis-related gene expression.

## Author contributions

DY, BB, TB, TT and DY designed and directed the project; AD, WQ and HH performed the experiments. SS, TT, SC, CW and DY analyzed data and drafted the article. AD, TB and DY wrote the manuscript. All authors reviewed the results and approved the final version of the manuscript.

